# Repressing Integrase attachment site operation with CRISPR-Cas9 in *E. coli*

**DOI:** 10.1101/110254

**Authors:** Andrey Shur, Richard M. Murray

**Affiliations:** Biology and Biological Engineering, California Institute of Technology, Pasadena, CA 91125.

## Abstract

Serine integrases are bacteriophage proteins responsible for integrating the phage genome into that of the host. Synthetic biologists have co-opted these proteins into useful tools for permanent DNA logic, utilizing their specific DNA recombination abilities to build synthetic cell differentiation and genetic memory systems. Each integrase has a specific pair of DNA sequences (attP/attB sites) that it recombines, but multiple identical sites can result in unpredictable recombination. We have developed a way to control integrase activity on identical attP/attB sites by using catalytically dead Cas9 (dCas9) as a programmable binding protein that can compete with integrase for binding to specific attachment sites. Utilizing a plasmid that contains two identical Bxb1 attP sites, integration can be repressed up to 8 fold at either one of the two attP sites when guide RNA and dCas9 are present. Guide RNA sequences that bind specifically to attB, or either of two attP sites, have been developed. Future goals are to utilize this technology to construct larger and more complex integrase logic circuits.

## Introduction

Serine integrases play a vital role in the bacteriophage life cycle by allowing phage DNA to be inserted and extracted from the host’s genome [1]–[3]. These enzymes are particularly interesting and useful to synthetic biologists because they are inherently directional. A serine integrase tetramer will catalyze a recombination event between two heterotypic DNA “attachment” sites (attP and attB) [4]. Without the presence of a cognate directionality factor, the integrase will not allow the reaction to proceed backwards, between the recombination products of those sites (attL and attR) [5]. Serine integrases are also notable for not requiring any host factors or additional proteins to catalyze recombination.

The directionality and sufficiency of these proteins make them very useful for synthetic biology. In the context of synthetic gene regulation, integrases allow an additional control layer to supplement the use of transcription factors in controlling gene expression [6]–[9]. Integrase attachment sites can be placed flanking a critical gene expression component such as a promoter or terminator or protein coding sequence. Upon induction of integrase, the component in question can be excised or flipped. Since the changes are wrought on the DNA itself, the integrase acts as a simple and leak-proof toggle switch.

Nested integrase attachment sites can be used to create a region of DNA that is guided through a series of DNA state changes by the expression of integrases in sequence [6], [9]-[11]. These state changes can be coupled back to the expression of other genes, thereby creating a synthetic differentiation pathway for bacteria. This type of logic allows fewer inducer species to be used for attaining more different DNA states. Likewise, the use of nested integrase logic can allow a bacterium to ‘record’ the chronological order of events.

Building logic using nested integrase attachment sites requires inducing recombination between pairs of sites at different times. A single integrase species will bind to and recombine with all cognate sites present, so to accomplish nested logic, orthogonal integrase species must be used. This puts a limit on the number of nested sites that can be utilized, as there are only about 13 orthogonal integrases that have been characterized [8]. To allow expansion of integrase based logic networks, full control of individual attachment site activity is needed. In this work we investigate whether catalytically inactive CRISPR-Cas9 can be used to repress integrase activity by preventing integrase from binding to an attachment site.

## Integrase activity in TX-TL

TX-TL is a cell-free rapid prototyping environment that allows proof of concept testing of genetic circuits [12]. Before proceeding to investigate integrase site repression in TX-TL, initial tests were perfomed to confirm that integrase activity is measureable. After 12 hours at 30°C, almost all plasmids in the reaction mixture containing attP and attB sites have been recombined (Figure 1). This demonstrates that integrases are produced and active in TX-TL.

**Figure 1:**
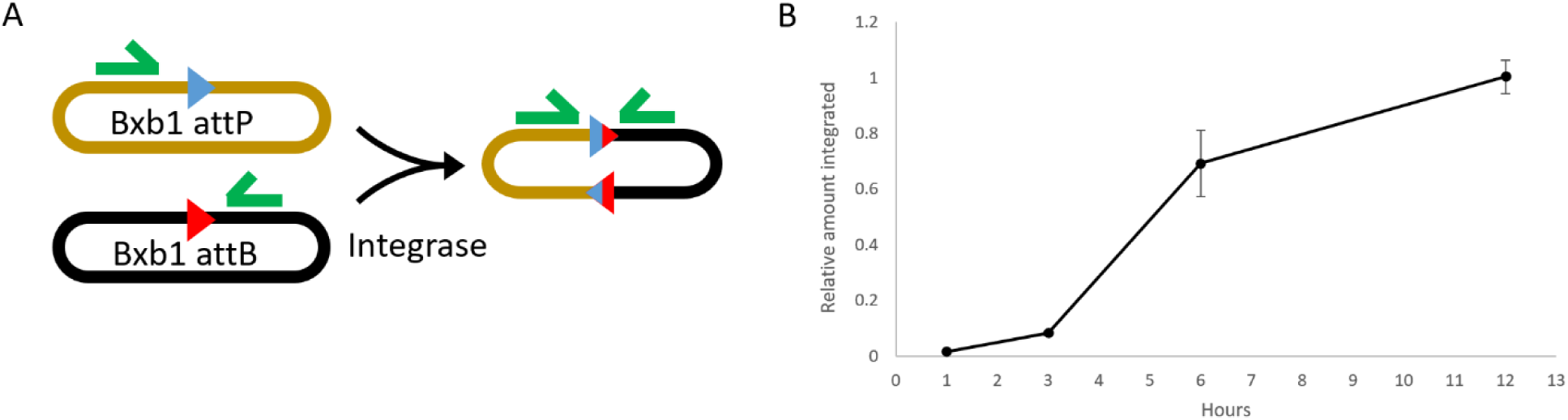
Time series of integration in TX-TL. Plasmid expressing Bxb 1 integrase and two plasmids containing Bxb1 attachment sites were incubated in TX-TL at 30°C, with samples taken at different times and qPCR was performed on the samples. (A) Blue and red triangles represent integrase attachment sites, green half-arrow shapes represent qPCR primers. Primers were designed to detect the DNA species formed following integration. (B) Time series data indicate that reaction reaches completion after 12 hours.

## dCas9 attachment site repression

Catalytically non-functional cas9 (dCas9) can be used as a programmable DNA binding protein [13]. Cas9 is known to bind strongly to DNA, and affect the function of other DNA-binding proteins such as RNA polymerase. We sought to use this binding activity to out-compete the binding of integrase to attachment sites specified by different guide RNAs. As well as needing a cognate gRNA, dCas9 requires the presence of a protospacer adjacent motif (PAM) sequence to the 3’ of the guide sequence [14]. In the case of SpCas9, this PAM is NGG. A test construct was made such that two identical attP sites are presented on the same vector, but only one of them has NGG in close proximity to the integrase attachment site. This allows a guide RNA to be designed that only binds to one of the two sites. When binding of dCas9 is activated by the production of guide RNA, integrase activity at one of two identical sites is decreased (Fig. 2). One interesting observation is that the more guide RNAs are being produced, the weaker the integrase activity repression. This may indicate that 1nM is already a saturating amount of guide RNA, and more guide plasmid simply consumes the resources of TX-TL faster, such that less dCas9 is produced.

**Figure 2:**
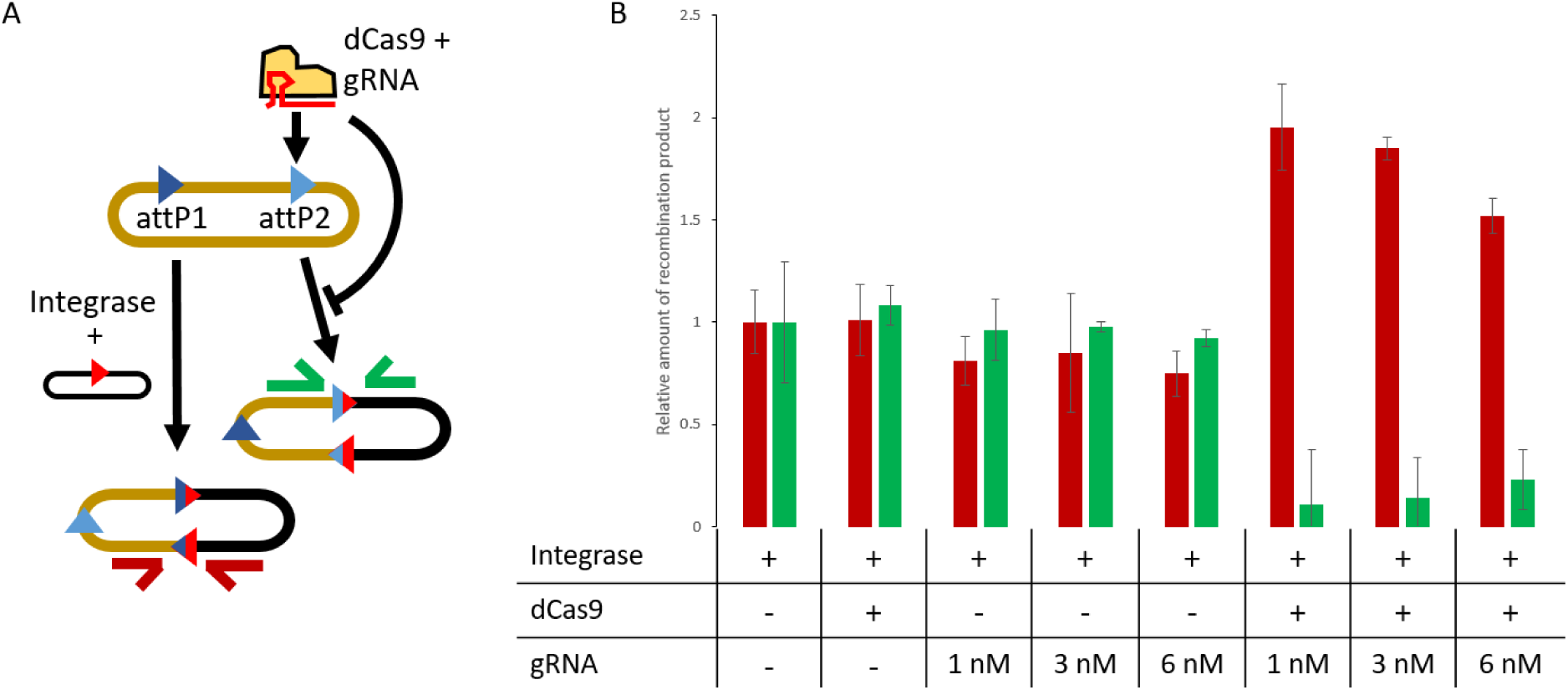
Integrase activity can be inhibited by the presence of dcas9 and a guide RNA designed to bind to one of two identical attP sites. (A) binding of dCas9 prevents integrase from binding and performing recombination at attP 2. (B) Expressed in TX-TL, integrase reacts at sites 1 and 2 at approximately the same rate if gRNA or cas9 are added independantly, but when both components are present, site 2 (green bars) is recombined significantly less often than site 1 (red bars).

## Additional guide RNA testing

It is difficult to predict the effectiveness of guide RNA sequences without empirical observations, so a variety of different guide RNAs were designed and tested (Figure 3). Some guide RNAs, such as L2 (the same one tested in Figure 2) and B1, are extremely effective at shutting down targeted site activity. Other guides, such as M1, M2, and L1, have lower effectiveness. The lower effectiveness of M1 and M2 when compared to B1 is difficult to explain. More guide testing must be done before any generalizations can be made about effective guide design practices for blocking integrase activity.

**Figure 3:**
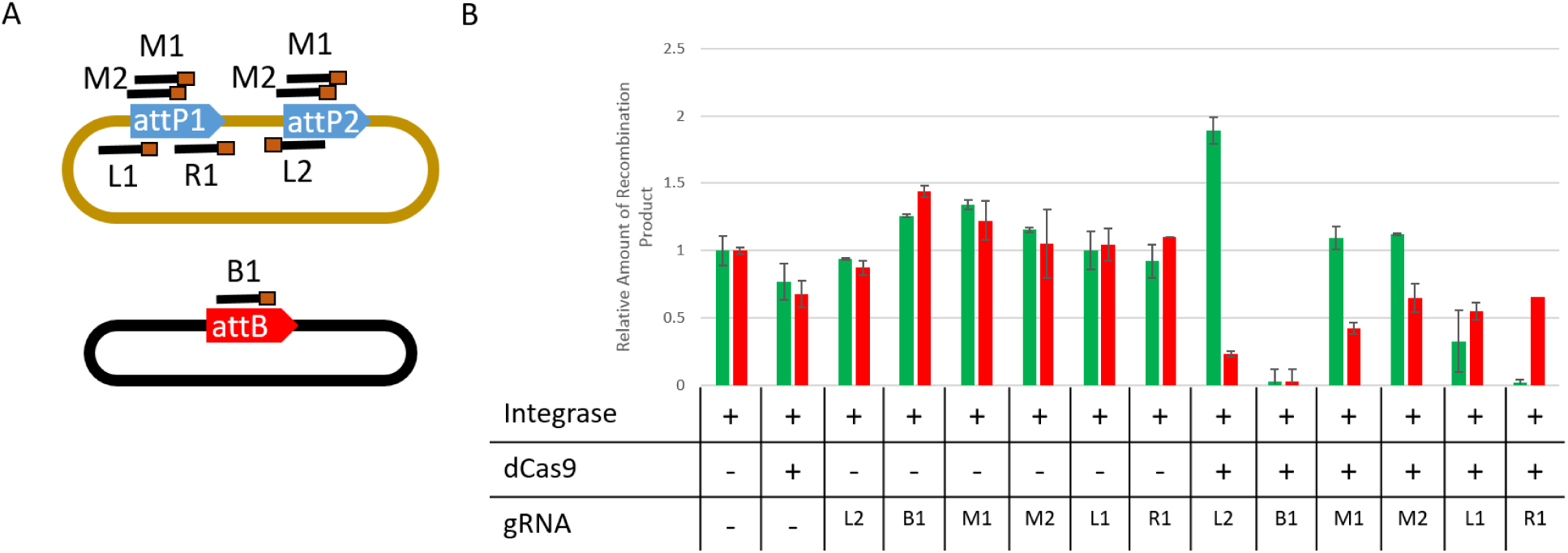
Testing of additional guide RNAs. (A) Map of guide RNA locations. Brown box represents the location of the NGG PAM sequence for each gRNA. M1 and M2 bind to both attP sites (B) repression effects from each guide. Red bars represent integration from site 1, and green bars represent integration from site 2.

## Conclusion

Results presented here are intended as a stepping stone to developing a new dCas9-based control layer for integrase logic circuits. Future work will focus on moving the dCas9 attachment site to cells and amassing a larger catalog of confirmed working guide RNA sequences. The goal of this work is to allow arbitrary control of integrase attachment site function with expression of guide RNAs, thus the dynamics and effectiveness of this method must be quantified. It may be that strategic positioning of the guide RNA binding site relative to the integrase attachment site leads to better repression, or that some integrases are not compatible with dCas9 repression. Another interesting question is how much the added stress of having to express dCas9 and guide RNAs adversely affects the performance of integrase-based logic circuits. There may also be some dCas9 fusion proteins that can make integrase site repression more or less efficient.

## Materials and Methods

TX-TL extract was produced and reactions performed as described previously [15]. Bxb1 integrase sequence was amplified from the Dual-recombinase-controller vector, which was a gift from Drew Endy (Addgene plasmid # 44456)[9]. pAN-PTet-dCas9 was a gift from Christopher Voigt (Addgene plasmid # 62244)[16]. Sequences are as follows: Site1:

ATTCAGTCGTCACTCATGGTTCGTGGTTTGTCTGGTCAACCACCGCGGTCTCAGTGGTGTACGGTACAAACCCAAGCTCCAACATGG, attB: TCGGCCGGCTTGTCGACGACGGCGGTCTCCGTCGTCAGGATCATCCGGGC, Site2: CCAAATGTCGTGGTTTGTCTGGTCAACCACCGCGGTCTCAGTGGTGTACGGTACAAACCCTGAG, L2: ACCAGACAAACCACGACATT, B1: GGCCGGCTTGTCGACGACGG, M1: GTTTGTCTGGTCAACCACCG, M2: GTCAACCACCGCGGTCTCAG, L1: CTCATGGTTCGTGGTTTGTC, R1: TACAAACCCAAGCTCCAACA

## Acknowledgements

AS was supported by the NIH training grant number 5T32GM007616-37. Research supported in part by the Institute for Collaborative Biotechnologies through grant W911NF-09-0001 from the U.S. Army Research Office. The content of the information does not necessarily reflect the position or the policy of the Government, and no official endorsement should be inferred

